# Independent directional tuning of the human triceps surae muscles during standing postural control

**DOI:** 10.1101/2025.06.24.660756

**Authors:** Martin Zaback, Christopher K. Thompson

**Affiliations:** Department of Health and Rehabilitation Sciences, Temple University, Philadelphia, PA, USA

**Keywords:** Soleus, Gastrocnemius, Posture, Balance, Muscle synergies, Electromyography, Directional tuning

## Abstract

The triceps surae, composed of the soleus (SOL) and medial (MG) and lateral (LG) gastrocnemii, are anatomically-derived synergists which act as a functional unit to plantarflex the ankle. However, anatomical differences suggest that each muscle is capable of generating distinct torques at the ankle, raising the possibility that each can be independently controlled to suit the needs of a given task. This possibility was explored by investigating the activation patterns of the triceps surae during two balance tasks that use different neuromechanical control strategies to maintain equilibrium.

High-density surface EMG was recorded from the triceps surae of 14 healthy young adults during multiple trials of dual-and single-legged standing. Newly developed analyses examined how each muscle tuned its activity with center of pressure (COP) movement throughout 2-D space. During dual-legged standing, only the SOL and MG were active and both tuned their activity uniformly with anteroposterior COP movement. By contrast, during single-legged standing, each muscle showed robust activation and significantly different directional tuning, with the LG most active before medial COP movement, while SOL and MG were most active before lateral COP movement. Further analyses demonstrated the LG could be activated entirely independent of the SOL and MG, and vice versa, with independent activation of each muscle causing different angular deflections of the COP during single-, but not dual-legged standing. These observations reveal a sophisticated level of neural control, whereby the nervous system exploits subtle differences between highly similar muscles to tune balance corrective adjustments in a task-dependent manner.

**Significance:** The triceps surae, composed of the soleus and medial and lateral gastrocnemii, are anatomical synergists with shared roles in plantarflexion. By applying new analytical techniques to high-density surface EMG and kinetic data, we show that each triceps surae muscle can produce directionally tuned torques at the ankle joint and the nervous system exploits this functional heterogeneity to regulate balance corrective adjustments in a task-dependent manner. Analyses of regionally-defined motor unit subpopulations revealed negligible intramuscular functional differences, suggesting that each triceps surae muscle are treated as homogenous, but independent, actuators. This fractionated control of closely related lower limb postural muscles is likely critical to the maintenance of equilibrium and may be impaired by musculoskeletal and neurological impairments.

## Introduction

The human upright posture is inherently unstable, as a relatively large mass is balanced high above a small base of support. During bipedal standing, the center of gravity sits slightly in front of the ankle joints, resulting in a gravitational torque which continually acts to pull the body forward (Winter, 2009). The intrinsic stiffness of the Achilles tendon and other passive tissues are insufficient to counteract this gravitational torque, meaning that additional, active plantar flexor torques are needed to maintain upright balance (Morasso and Schieppati, 1999; Loram et al. 2002). In humans, the principal plantar flexors are the triceps surae, consisting of the soleus (SOL) and medial (MG) and lateral (LG) heads of the gastrocnemius. Despite sharing the same distal tendon, each muscle is remarkably distinct, differing in terms of their muscle fibre composition (Johnson et al. 1973; Edgerton et al. 1975), response to different modes of afferent input (Duysens et al. 1996; Dakin et al. 2016; Mildren et al. 2019), and pattern of activation while standing (Heroux et al. 2014; Vieira et al. 2012). Critically, there are also anatomical differences, with each muscle not only having separate origins, but also partitioned attachments onto the calcaneal tuberosity via distinct subtendons (Edama et al. 2016). Thus, while the triceps surae are ostensibly a functional unit, there are multiple physiological and anatomical differences which suggest they could have independent roles during balance control.

Of particular interest are the potential independent roles of the triceps surae in generating balance corrective torques outside of the sagittal plane. This is typically overlooked since static balance is often studied during dual-legged standing – a posture where mediolateral sway is controlled primarily by loading and unloading weight between the hips rather than by inverting and everting the ankles (Day et al. 1993; Winter et al. 1993; 2003). However, work that has mechanically stimulated each muscle of the triceps surae in human cadavers has demonstrated that each muscle is capable of generating distinct torques along the frontal plane, with significant eversion torques generated by LG and inversion torques generated by SOL and MG (Arndt et al. 1999). Since recent work suggests that each muscle of the triceps surae receives largely independent neural drive, as demonstrated by low or absent intermuscular coherence (Hug et al. 2021; Rossato et al. 2022) and individuals’ capability to volitionally modulate MG and LG motor units independent of one another (Rossato et al. 2024), it is possible that the nervous system can flexibly exploit the unique physiological and anatomical features of each muscle depending on the needs of a given balance task.

This possibility was explored in the present study by investigating the activation patterns of the triceps surae during two balance tasks that use different neuromechanical control strategies to regulate mediolateral stability: single-and dual-legged standing. The primary analyses of this study examined muscle activity and motor unit discharge time-locked to prominent center of pressure (COP) peaks occurring throughout 2-dimensional space. This was done to determine if the nervous system independently tunes neural drive to each muscle of the triceps surae depending on the task and orientation of the balance corrective adjustment. It was hypothesized each muscle would demonstrate uniform directional tuning along the sagittal plane during dual-legged standing, but highly divergent tuning during single-legged standing - a task where ankle inversion/eversion is needed to regulate equilibrium. Additional analyses investigated if directional tuning is further refined within each muscle through distinct motor unit task groups (Loeb 1985; Segal et al. 1991). Through these and other confirmatory analyses, we demonstrate that each muscle of the triceps surae generates vastly different torques at the ankle joint and the nervous system exploits these functional differences to tune balance corrective adjustments in a task-dependent manner.

## Methods

### Participants

Fourteen healthy young adults (10 male; mean age: 29.9±7.2 years) participated in this study. Participants were free of neurological or musculoskeletal disorders that could impair their balance or movement. All procedures were cleared by Temple University’s Institutional Research Ethics Board and complied with the Declaration of Helsinki, except for registration in a database. Participants provided written informed consent.

### Experimental design

Participants stood on two independent force plates (Fully Instrumented Treadmill, Bertec, OH, USA) and completed 8 trials of single-and dual-legged standing (16 trials total). Dual-legged trials were each approximately 2 minutes and participants stood with a stance width equal to the length of their foot, with each foot positioned on separate force plates. The angle of each foot was self-selected, although no more than 5° of lateral rotation was permitted for either foot. Single-legged trials were approximately 30-s and participants stood on their right leg. The position of participants’ feet were traced onto the support surface to ensure consistent foot orientation was maintained over the course of the experiment. During each trial, participants were instructed to stand quietly with their arms resting at their sides while maintaining visual focus on an eye-level fixation point approximately 2.5 meters away. Trials were completed in a blocked order, with all dual-legged trials completed first to minimize fatigue associated with single-legged standing influencing this condition. Participants wore a safety harness that was mounted to an overhead track. The tension of the harness was adjusted such that support was only provided in the event of a fall. In addition, hand rails were positioned along both sides of the participant. One participant (male) consistently required support during each single-legged standing trial, either grabbing the handrails or stepping down multiple times with their non-support leg; thus, this participant’s single-legged standing data were excluded from analyses.

### Surface EMG and recordings

The skin over the right SOL, MG, and LG were shaved, lightly abraded, and cleansed with water in preparation for the placement of 64-channel high-density electrode arrays. The electrode consisted of a 5×13 electrode matrix with 8 mm inter-electrode distances (GR08MM1305, OT Bioelettronica, Torino, Italy). Electrode placement was guided by palpation and visible borders of the triceps surae seen during plantarflexion contractions. The electrodes over the MG and LG were oriented vertically over the center of each muscle belly, with the lower margin of the electrode located ∼1-2 cm from the most distal portion of each muscle. The SOL array was placed ∼1-2 cm below the distal portion of the medial gastrocnemius muscle, oriented vertically and medial to the midline of the Achilles tendon. In a subset of participants (n=9), a second array was placed over the lateral portion of the SOL. Only data from the medial SOL array (henceforth referred to as the SOL array) were included in the main analyses for this study, as there was significant contamination from putative fibularis motor units across a substantial portion of the lateral SOL array. A wetted strap electrode was secured around the right ankle and was used as a ground. Differential EMG data were analog filtered (10-900 Hz), amplified (150x), and sampled at 2048 Hz using a 16-bit A-D converter (Quattrocento, OT Bioelettronica, Torino, Italy).

### Force plate recordings

Forces and moments from each force plates were amplified and sampled at 1000 Hz using a separate 16-bit A-D converter (Lock Lab, Vicon Motion Systems, UK). In order to synchronize EMG and force plate recordings, a single channel of SOL EMG was converted to an analog output and digitized by both A-D converters during all trials. A customized cross-correlation algorithm (Matlab, MathWorks, MA, USA) was used offline to determine the precise time-delay and relative sampling rate between the separate recordings (Zaback et al. 2022). Force plate data were then resampled to 2048 Hz and time-aligned with the EMG.

### Surface EMG preprocessing

Recordings from high-density electrode arrays were visually inspected offline and channels containing visible artifacts were manually excluded from further analyses.

Unless stated otherwise, analyses of surface EMG interference signal were limited to a single, artifact-free channel of differential EMG from the center of each array. The same channel was examined across all trials within a given subject. This minimized cross-talk with adjacent muscles and facilitated comparison of muscle activation patterns between tasks. EMG data were high-pass filtered offline (20 Hz cut-off, 2^nd^ order dual-pass Butterworth filter) and rectified prior to subsequent analyses.

### Motor unit decomposition

All artifact-free EMG channels from each electrode array (maximum of 63) were decomposed into the spike-times of single motor units using a convolutive blind source separation algorithm (Negro et al. 2016) with a silhouette threshold set to 0.87. To improve decomposition accuracy, an experienced investigator used a semiautomated motor unit cleaning algorithm to manually edit non-physiological motor unit spike times. Specifically, the automatic decomposition results were updated iteratively by re-estimating each motor unit spike train after correcting for missing spikes or spikes that deviated substantially from the discharge profile (Del Vecchio et al. 2020).

### Force plate preprocessing

Forces and moments from both force plates were low-pass filtered offline (5 Hz cut-off, 2^nd^ order dual-pass Butterworth filter) and the COP under each foot was calculated along X (mediolateral) and Y (anteroposterior) axes. For single-legged standing, COP from the right force plate was analyzed, while for the dual-legged condition, the net COP was analyzed as the weighted average of both right and left COP signals after both had been aligned to the same global coordinate system. This was done because, unlike during single-legged standing, the COP underneath each foot during dual-legged standing generally moves only along the sagittal plane, with negligible movement along the frontal plane (Winter et al. 1993).

### Tuning curve analyses – surface EMG

Previous work has shown that there are phasic changes in muscle activity time-locked to discrete reversals in COP position (Loram et al. 2005; Zaback et al. 2023). For example, there is a phasic increase in SOL EMG and concurrent decrease in tibialis anterior EMG which reach local maxima and minima, respectively, ∼150 ms before anterior peaks in COP position (Loram et al. 2005; Zaback et al. 2023). Thus, examination of muscle activity time-locked to discrete reversals in COP position can provide insight into the phasic changes in neural drive to different motor pools.

A rotation matrix was applied to the 2-D COP data to permit identification of prominent COP peaks occurring in different directions across 2-D space. Rotations were applied iteratively in 5° clockwise increments, creating a total of 72 unidimensional, rotated COP time-series (COP_r_). With this convention, positive peaks in the COP_r_ time-series are purely forward at 0° of rotation, right at 90° of rotation, backward at 180° of rotation, and left at 270° degrees of rotation. While the profile of COP_r_ time-series only changes slightly with each iterative rotation, clear changes in the timing of identified COP_r_ events can be seen with coarse changes in rotation (Figure 1CD).

**Figure 1.**
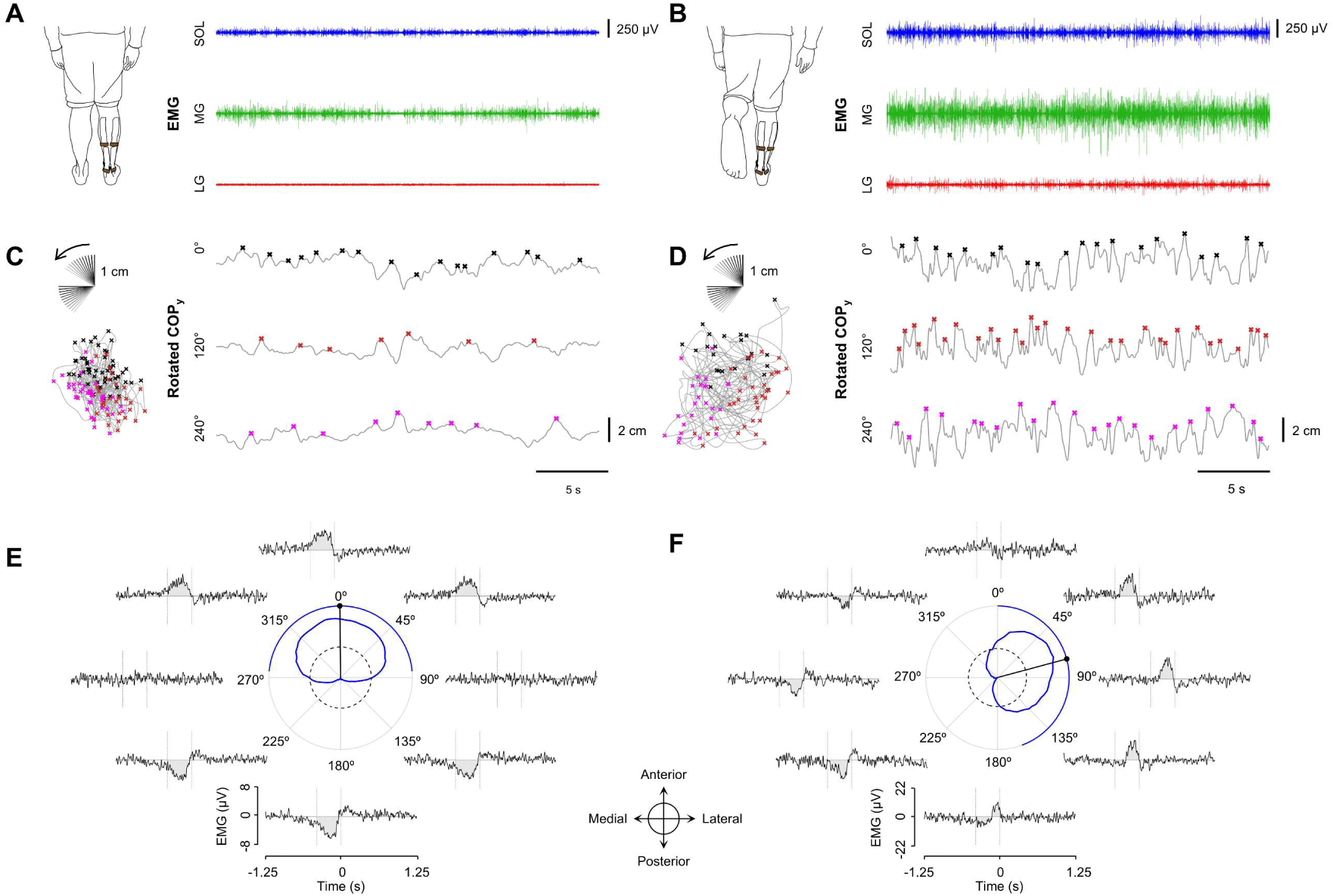
Experimental set-up and EMG tuning curve analysis. High-density surface EMG was recorded from the medial (MG) and lateral (LG) heads of the gastrocnemii and medial and lateral aspects of the soleus (SOL) while participants performed trials of dual-and single-legged standing on independent force plates (A, B). Prominent positive peaks from the COP_y_ time-series were identified iteratively as the COP coordinate system was rotated counterclockwise in 5° increments. A subset of rotated unidimensional COP time-series from representative trials of dual-(C) and single-legged (D) standing are illustrated with corresponding COP events indicated with x’s. Insets of the original COP_xy_ data are shown next to each rotated COP time-series with corresponding events shown in 2-D space (color-coded x’s). From each rotated unidimensional COP time-series, rectified EMG were trigger-averaged from corresponding COP events (E, F), creating a series of 72 waveform averages for both stance conditions. EMG tuning curves were then created by calculating area under the curve of each waveform average between-400ms – 0 ms relative to each event (shaded area on each EMG waveform average). Representative EMG tuning curves from the SOL are shown in the center of E and F from a representative participant during dual-and single-legged standing, respectively. The dashed inner circle corresponds to an area under the curve value of 0, with data below representing inhibition of muscle activity preceding COP peaks in a given direction, and data above representing excitation. The circular mean of each tuning curve is shown as a single black line capped by a filled circle, while a radial band represents the region where area under the curve values were greater than 20% of the maximum excitatory response. Tuning curves are oriented such that 0° corresponds with the magnitude of EMG preceding pure forward COP peaks, while 90° corresponds with the magnitude of EMG preceding pure rightward (lateral) COP peaks.

COP_r_ events were identified as positive peaks in the COP_r_ time-series that exceeded a threshold based on the variability of COP_r_ oscillations within each trial. This threshold was calculated as 1 standard deviation (SD) of the high-pass filtered (0.1 Hz cut-off, 2^nd^ order dual-pass Butterworth) COP_r_ time-series (Zaback et al. 2023). For an event to be identified, a peak in the COP_r_ time-series had to exceed the 1 SD threshold in terms of its prominence (height relative to adjacent peaks; Figure 1CD).

After COP_r_ event times were calculated across all trials for each iteration of rotation, EMG waveform averages were created from the rectified EMG separately for both tasks. This was done by averaging EMG around a ±1.5 s window centered on each event time across all rotations. This resulted in a total of 72 unique waveform averages for each muscle during each task. Each waveform average was baseline corrected by subtracting the mean EMG from-1500 ms to-1000 ms relative to the event time. The magnitude of each muscle’s response was then calculated as the area under the curve (AUC) from-400 ms to 0 ms (Figure 1EF). Once this operation was completed for all 72 waveform averages, an EMG tuning curve was created by plotting AUC values across polar coordinates, with each AUC value indicated by a radial distance at an angle corresponding to the rotation that had been applied to the COP to create the respective waveform average from which the AUC value was derived (Figure 1EF). From these data, the circular mean of the distribution was calculated (Berens, 2009) which represents the angular orientation of COP movement most often associated with the greatest excitatory drive to each muscle. In addition, to illustrate the directions of COP movement where each muscle received significant excitatory input, regions where AUC values exceeded 20% of the maximal AUC were calculated and plotted as a radial band outside of each tuning curve (Figure 1EF).

A surrogate of this analysis was repeated where COP_r_ event times were shuffled, such that the mean and variance of the events were preserved, but were no longer systematically related to true peaks in the COP_r_ signal. This was done to confirm that this analysis does not inherently generate non-uniform circular distributions, but rather only does so when muscles demonstrate true directional tuning with balance corrective torques (Figure S1).

Additionally, to confirm electrode location did not significantly impact estimates of directional tuning, EMG tuning curves were constructed for all channels of differential EMG spanning the electrode array for each muscle (Figure S2). Five of the 63 possible differential channels were not included in the construction of the EMG tuning maps since these differential channels were calculated from horizontally adjacent, as opposed to vertically adjacent pairs of electrodes. All remaining channels were included so long as their recordings were consistently artifact-free. The circular means from each tuning curve spanning a single array were calculated and pooled together so that a circular standard deviation could be calculated to provide an estimate of variance in directional tuning across the surface of each muscle (Berens, 2009).

### Motor unit discharge characteristics

Mean discharge rate and coefficient of variation (CoV) of discharge rate were calculated for each motor unit during periods of repetitive discharge (i.e., interspike intervals (ISIs) <400 ms). An intermittency index, adapted from Cohen and colleagues (2023), was calculated to characterize the extent of recruitment and derecruitment of each motor unit over time. This index was calculated as the number of times a motor unit was recruited (defined here as discharge of at least 2 spikes with ISI <400 ms) and derecruited (defined as an increase in ISI >400 ms) per minute.

### Tuning curve analyses – motor units

Using the same COP_r_ event time-stamps detailed above, peristimulus-time histograms (PSTHs) and peristimulus frequencygrams (PSFs) were constructed from motor unit spike times. PSTHs provide an estimate of the probability that a motor unit will discharge within a given period of time relative to a specific event. In the current study, these were created by tallying the number of spike times occurring within 20 ms bins across a ±1.5 s peristimulus period centered around each COP_r_ event. Spikes were counted cumulatively across all motor units within each muscle for all trials of the same task separately for each iteration of COP rotation. PSFs quantify the change in discharge rate of repetitively firing motor units within a given period of time relative to a specific event. Unlike probability-based measures (i.e., PSTHs and waveform-averaged EMG), PSFs are less influenced by the discharge history of a motor unit and provide a more accurate estimate of changes in the net synaptic input to motoneurons (Turker and Powers, 2005). To create PSFs, the instantaneous discharge rate of each motor unit was calculated as the inverse of the ISI. These values were then superimposed relative to each COP_r_ event time over a ±1.5 s peristimulus period for each muscle, only including those motor units that were repetitively firing throughout the pre-stimulus period (>3 spikes with ISIs<0.4 s). A total of 72 unique composite PSTHs and PSFs were generated for each muscle, each representing changes in motor unit firing probability and rate coincident with COP reversals in different directions, respectively.

Unless stated otherwise, analyses focused on tuning curves constructed from PSFs since they provide a physiologically validated estimate of the underlying synaptic input to motoneurons (Turker and Powers, 2005). PSF tuning curves were generated by calculating the baseline-corrected (-1500 to-1000 ms) discharge rate (ΔDR) within two 400 ms windows before (-350 to 50 ms) and after (50 to 450 ms) the peak of COP_r_ reversals. The second window was selected to analyze an apparent secondary synaptic event of opposite polarity that was consistently observed after the peak of COP_r_ reversals. PSFs were essential for this particular part of the analysis to determine if this putative secondary synaptic event truly represented a distinct change in motoneuron postsynaptic potentials, or was instead an epiphenomenon caused by phase-delayed or phase-advanced spikes resulting from the initial synaptic event (Turker and Powers, 2005). Once ΔDR values for both the initial and secondary synaptic events were calculated across all 72 iterations of COP rotation, tuning curves were constructed using the same procedures described previously and circular means were calculated from each distribution (Berens, 2009). When tuning curves were constructed from PSTHs, the same general procedures were used, with the exception that peristimulus windows for the initial and secondary synaptic events were 400 ms immediately before (-400 to 0 ms) and after (0 to 400 ms) each COP reversal. Tuning curves for a given muscle were only constructed if, on average, at least 1 sufficiently long motor unit spike train (>100 spike for single-legged stance; >300 spikes for dual-legged stance) could be decomposed across respective trials for each condition (i.e., at least 8 sufficiently long motor unit spike trains total within a given muscle and condition).

### EMG burst analyses and 2-D COP trajectories

To determine if muscles of the triceps surae can be activated independently of one another to generate distinct balance corrective torques, EMG and COP data were time-locked to isolated bursts of activity from each muscle of the triceps surae. Smoothed EMG signals were created by low-pass filtering (2 Hz low-pass, 5^th^ order, dual-pass Butterworth filter) the rectified EMG from each muscle and then the same peak detection algorithm described previously was used to identify prominent increases in EMG. These smoothed EMG event times were used to create waveform averages (±1.5 s) of rectified SOL, MG, and LG EMG (EMG-EMG) and COP_x_ and COP_y_ (EMG-COP).

All waveform averages were baseline-corrected by subtracting the mean from-1500 to - 1000 ms relative to each EMG event time.

To examine if muscles of the triceps surae could be activated independently of each other, mean activity of each baseline-corrected EMG-EMG waveform average was calculated between a ±250 ms period centered on each EMG burst event. To determine if each muscle generated distinct balance corrective torques, COP_x_ and COP_y_ waveform averages for each event-type were combined, creating a 2-D COP trajectory. The apex of the 2-D COP trajectory was identified as the farthest point from the baseline position occurring within 300 ms after the EMG bursts. The distance and angle of the resultant vector between the origin and apex was calculated, providing an estimate of the magnitude and direction of COP movement associated with isolated activation of each muscle.

### Regional tuning curve analyses – motor unit subpopulations

To determine if regionally confined motor unit subpopulations within each muscle have distinct directional tuning, PSTH tuning curves were constructed for groups of motor units located across the extreme margins of each array. These regional analyses were only performed on data from the single-legged stance condition, as EMG and motor unit tuning curves at the level of the whole muscle were highly heterogeneous during this task only. Motor unit locations were first determined by spike-triggered averaging surface EMG across all differential recording sites. This reconstructed the profile of each motor unit action potential (MUAP) seen across the surface of the array. From these data, the peak-to-peak amplitude of the MUAP at each site was calculated. The centroid was determined as the average X and Y coordinates of all recording sites where the peak-to-peak amplitude of the MUAP was greater than 90% of the largest MUAP. Motor units from each muscle were then classified according to the location of their centroid.

Motor units located in the upper and lower quartiles along the X-axis were classified as lateral and medial units, respectively. Motor units located in the top and bottom quartiles along the Y-axis were classified as proximal and distal units, respectively. After all units from each muscle were classified across all trials for each participant, PSTH tuning curves were constructed separately for each motor unit classification using the same procedures described previously. For each possible within muscle comparisons (i.e., medial vs lateral and proximal vs distal), a participant’s data was only included if each PSTH tuning curve was constructed from 4 or more sufficiently long motor unit spike trains (> 100 spikes) and there was significant separation between the average motor unit centroid location within each group (≥15 mm along the x-axis for medial vs lateral comparisons and ≥40 mm along the y-axis for proximal vs distal comparisons).

### Regional tuning curve analyses – medial and lateral SOL

PSTH tuning curves were constructed separately for motor units from medial and lateral array in 9 participants (5 males) where separate hd-EMG recordings were made from medial and lateral portions of the SOL. Prior to performing these regional analyses, EMG tuning curves were constructed for all channels of differential EMG spanning both electrode arrays. These analyses revealed that recordings made from the distal and lateral corner of the lateral SOL array were consistently tuned in the opposite direction as all other SOL EMG channels, suggesting the presence of cross-talk from adjacent fibularis muscles or stark functional differences. Therefore, motor units decomposed from the lateral SOL array for were stratified based on their centroid location, with those located in the distal corner categorized as putative fibularis and those from the remainder of the array as true lateral SOL. The boundary demarcating this stratification was set according to the region where differential EMG channels from this array were diametrically opposed to the rest of the SOL channels across all participants. Motor unit characteristic as well as the peak-to-peak amplitude of waveform-average MUAPs were calculated for each unit classification. As putative fibularis motor units consistently displayed highly intermittent firing with relatively few spikes per spike train, a more liberal criteria for including individual participant fibularis motor unit data in the construction of PSTH tuning curves was permitted (≥4 spike trains with >40 spikes across all trials of single-legged standing).

In a separate experiment aimed to investigate possible mediolateral gradation of SOL tuning curves, one male participant (33 years old) completed 10 trials of single-legged standing while intramuscular EMG recordings were made from pairs of electrodes spanning the mediolateral margins of the SOL. As with the main experimental protocol, trials lasted approximately 30-s and were completed while standing on a force plate.

Each fine wire electrode consisted of a sterilized pair of perpendicularly cut, 0.75 mm-diameter insulated stainless steel wires, that were inserted into the muscle via a 25-gauge hypodermic needle (Spes medica, Genova, IT). A total of 6 pairs of intramuscular electrodes were inserted into the SOL. Insertions were made 2 cm below the distal margins of the gastrocnemii, and spanned the mediolateral margins of the SOL at 2 cm increments. The third and fourth pairs of electrodes straddled the midline of the SOL, such that the third pair was 1 cm medial of the midline, while the fourth pair was 1 cm lateral. Intramuscular recordings were bandpass filtered (100-4400 Hz) and amplified (150x), and sampled at 10,240 Hz using the same16-bit A-D converter (Quattrocento, OT Bioelettronica, Torino, Italy). EMG tuning curves were constructed from each rectified intramuscular recording using the same procedures described previously.

### Statistical analysis

Watson-Williams (WW) tests, which examine if the average angular direction of two or more samples significantly differ (Berens, 2009), were used to compare the orientation of tuning curves separately across tasks, muscles, and motor unit subpopulations.

Specifically, separate WW tests were used to compare the orientation of EMG tuning curve circular means between tasks within same muscle (task-dependent differences in tuning) and between muscles within the same task (intermuscular differences in tuning). WW tests were also used to compare PSF tuning curve circular means separately for initial and secondary synaptic events as well as for PSTH tuning curves circular means for regionally-defined motor unit subpopulations within muscles. As a number of participants had an insufficient yield of motor units for some muscles, the sample size of pairwise WW tests varied depending on the comparisons being made. A Monte Carlo simulation-based analysis was used to determine the smallest sample needed for sufficiently powered WW tests. This power analysis determined the incidence of statistically significant differences between angular values randomly sampled from two von Miser distributions that differed by 30° with concentration parameters of 15°. These parameters were selected because 1) the mean difference reflects a modest, but functionally meaningful difference in directional tuning between two muscles with similar, but distinct actions and 2) the concentration parameters were representative of the average dispersion of EMG tuning curves based on pilot data. Across 100000 simulations with these parameters, the rate of statistically significant effects was ∼87% for samples when at least 6 angular values per group. This means that comparisons with 6 or more samples per group would be sufficiently powered to detect meaningful effects and smaller sampled comparisons should be interpreted with caution.

Linear mixed effects models (LME) were used to examine how motor unit discharge characteristics (discharge rate, CoV of discharge rate, intermittency index) differed between muscles and tasks. Since motor units from the LG could not be consistently decomposed during dual-legged standing, the first model examined Muscle × Task interactions and main effects of Muscle and Task including only SOL and MG motor units. The second model examined main effect of Muscle during the single-legged stance condition for all muscles, as a sufficient yield of motor units was available for all three muscles during only this task. For each model, discharge characteristics from individual motor units were included as the dependent outcomes, while participant identification number was treated as a random effect. To avoid including the same motor unit multiple times in the same model, comparisons were made with data from a single trial from each condition that had a high motor unit yield from each muscle. Significant interactions were followed up with Bonferroni-corrected pairwise comparisons that examined the effect of Muscle separately across both levels of Task and the effect of Task across both levels of Muscle. Significant main effects of Muscle were followed up with Bonferroni-corrected pairwise comparisons; follow-up comparisons for significant effects of Task were not necessary since this fixed factor only had two levels.

Single-samples t-tests examined if mean activity of each EMG-EMG waveform average were significantly different from zero. Non-significant effects indicate that phasic increases in activity of one muscle are not accompanied by significant modulation of another, whereas significant effects below or above zero indicates phasic of activity of one muscle is accompanied by significant inhibition or coactivation of the other, respectively. WW tests were used to compare the angular orientation of 2-D COP trajectories time-locked to isolated bursts of each muscle during both dual-and single-legged tasks. One-way repeated measure ANOVAs were used to compare the resultant distance of 2-D COP trajectories time-locked to isolated bursts of each muscle during dual-and single-legged tasks. Significant main effects of Muscle were followed up with Bonferroni-corrected pairwise comparisons.

All circular statistics were performed using the CircStat toolbox in Matlab (Berens, 2009), while all other statistical analyses were performed using RStudio (Posit PBC, Boston, MA, USA). For all comparisons, alpha was set to 0.05.

## Results

### Triceps surae demonstrate heterogenous and task-specific tuning of muscle activity while standing

During dual-legged standing, the SOL and MG modulated their activity with COP oscillations oriented primarily along the sagittal plane, such that each demonstrated increased activity before forward COP peaks and decreased activity before backward COP peaks (Figure 2AC). As the LG was generally quiescent during dual-legged standing (Figure 1A), analyses of EMG tuning curves were limited to the SOL and MG during this task. WW tests revealed that the circular means of SOL and MG tuning curves were significantly different (F_(1,26)_=4.2689, p=0.0489), with MG tuning curves oriented more laterally than the SOL (Table S1; Figure 2C).

**Figure 2.**
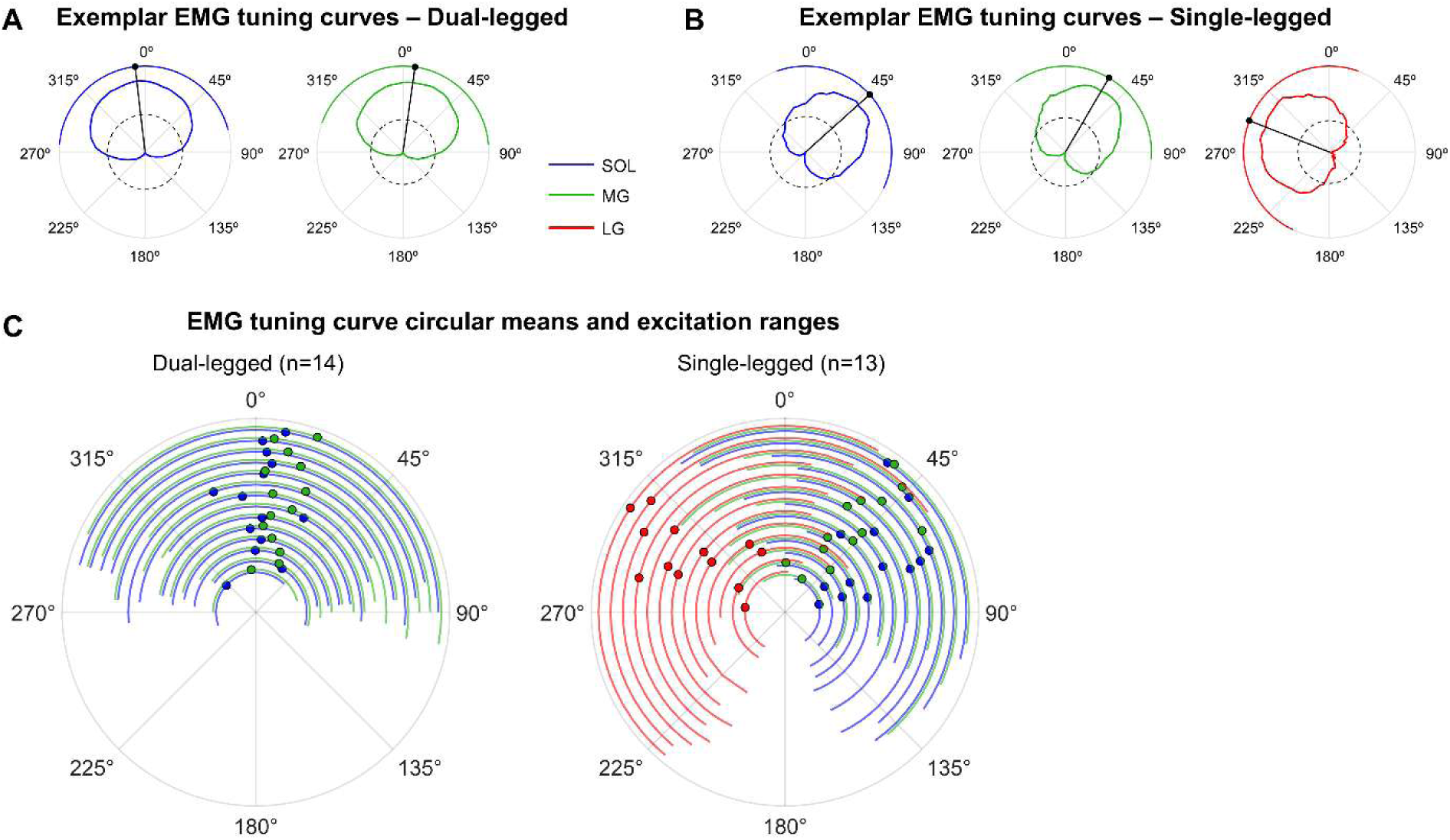
EMG tuning curve results. Exemplar tuning curves are shown for a single participant during dual-(A) and single-legged (B) standing conditions. Tuning curves were not analyzed for the LG during dual-legged stance condition since it showed little or no activity during this stance condition across all participants. (C) Circular means (●) from EMG tuning curves across all participants during dual-(n=14) and single-legged (n=13) standing conditions are shown along with radial-bands indicating the angles where area under the curve values for waveform averaged EMG were greater than 20% of the max excitatory response. Individual participant data are presented in radial steps, with data from each muscle of a single participant grouped together before incrementing outward to the next participant. *One participant’s data was excluded from the single-legged stance condition due to an inability to successfully perform the task.

By contrast, remarkably heterogenous EMG tuning curves were observed across all three muscles of the triceps surae during single-legged standing (F_(2,36)_=187.2114, p<0.0001). In particular, tuning curves for the SOL and MG were rotated significantly clockwise relative to their orientation during dual-legged standing (F_(1,24)_= 75.9662; p<0.0001; F_(1,24)_= 25.7077, p<0.0001, respectively), indicating that each muscle now had a significantly greater frontal plane component and increased their activity the most prior to forward and *lateral* COP movement. Unlike during dual-legged standing, the LG was highly active during single-legged standing (Figure 1B), permitting the analysis of EMG tuning curves. Tuning curves for the LG were approximately orthogonal to the SOL and MG, indicating it was more active prior to forward and *medial* COP movement (Figure 2C). WW tests confirmed that the circular means for each muscles’ tuning curves were significantly different, with LG tuning curves oriented more medially compared to SOL (F_(1,24)_=310.0426, p<0.0001) and MG (F_(1,24)_=217.8940, p<0.0001).

Furthermore, in contrast to what was observed during dual-legged standing, SOL tuning curves were oriented significantly more laterally compared to the MG (F_(1,24)_= 19.1899, p=0.0002).

### Motor units demonstrate similar directional tuning and reveal time-dependent antagonism between muscles of the triceps surae

A sufficient yield of SOL and MG motor units could be decomposed across both tasks for nearly all participants, whereas 7 participants (all male) yielded a sufficiently high number of LG motor units during the single-legged standing condition (Table S1).

Consistent with the low levels of the LG EMG during dual-legged stance, motor units could rarely be decomposed from the LG during this task.

Figure 3A and Table S1 provides the discharge characteristics of each muscle during dual-and single-legged standing. When considering motor units that could be compared across tasks (SOL and MG), significant Muscle × Task interactions were observed for all motor unit characteristics (discharge rate: F_(1,620)_ = 26.591, p<0.0001; CoV: F_(1,620)_ = 72.719, p<0.0001; intermittency: F_(1,626)_ = 106.536, p<0.0001). During dual-legged standing, SOL motor units were tonically active (as indicated by a low intermittency index) with low discharge rate variability (as indicated by a low CoV). This is in contrast to MG units, which demonstrated a higher (t_(621)_=4.979; p<0.0001) and more variable discharge rate (t_(622)_=8.960; p<0.0001) and more intermittent recruitment (t_(628)_=9.597; p<0.0001; Figure 3A). When performing single-legged standing, the firing characteristics of both muscles changed in distinct manners. While both muscles showed increases in mean discharge rate and CoV, increases in mean discharge rate were greater for the MG (t_(627)_=11.982, p<0.0001) than the SOL (t_(619)_=4.894, p<0.0001), while increases in discharge rate variability were greater for the SOL (t_(620)_=15.145, p<0.0001) than the MG (t_(626)_=3.104, p=0.0107). Furthermore, both muscles showed opposite changes in intermittency, with SOL becoming more intermittent (t_(626)_=6.950, p<0.0001) and MG becoming more tonic (t_(623)_=-7.457, p<0.0001) during single-compared to dual-legged standing. When contrasting the discharge behaviour of all 3 muscles of the triceps surae during single-legged standing, the SOL demonstrated significantly lower firing rates compared with both MG (t_(407)_=-11.351, p<0.0001) and LG (t_(409)_=-6.494, p<0.0001), while no significant differences in discharge rate were seen between MG and LG (t_(406)_=1.465, p=0.3089). LG units also demonstrated significantly greater discharge rate variability than both SOL (t_(406)_=3.234, p=0.0038) and MG (t_(403)_=4.913, p=0.0038), while both SOL (t_(409)_=4.214, p=0.0001) and LG (t_(410)_=4.237, p=0.0001) units were more intermittently recruited than MG units (Figure 3A), but did not differ from each other in term of intermittency (t_(411)_=1.229, p=0.4368).

From these motor unit data, composite PSFs were generated for each muscle using the same rotated COP events described in the initial analyses. Figure 3B illustrates individual participant PSFs for each muscle derived from COP events at three contrasting angles (230°, 350°, and 70°). These exemplar data demonstrate that at certain points in time, depending on the orientation of the balance corrective torque, the muscles of the triceps surae can receive the same or opposite synaptic inputs. For example, in the first column of Figure 3A, the PSF for the LG is of opposite polarity relative to the SOL the MG, indicating that as the COP moves medially and slightly posterior, the LG receives excitatory synaptic input (evident from a time-dependent increase in firing rate; Turker and Powers, 2005) prior to the peak of the COP, while, at the same time, the SOL and MG receive inhibitory inputs (evident from a concurrent decrease in firing rate). This same time-dependent antagonism is observed after the COP has peaked, as the LG receives inhibitory input while the SOL and MG receive excitatory inputs. This is in contrast to when the COP moves near purely forward, in which case each muscle of the triceps surae receive the same polarity of synaptic input before and after the peak of the COP (Figure 3B, middle column).

**Figure 3.**
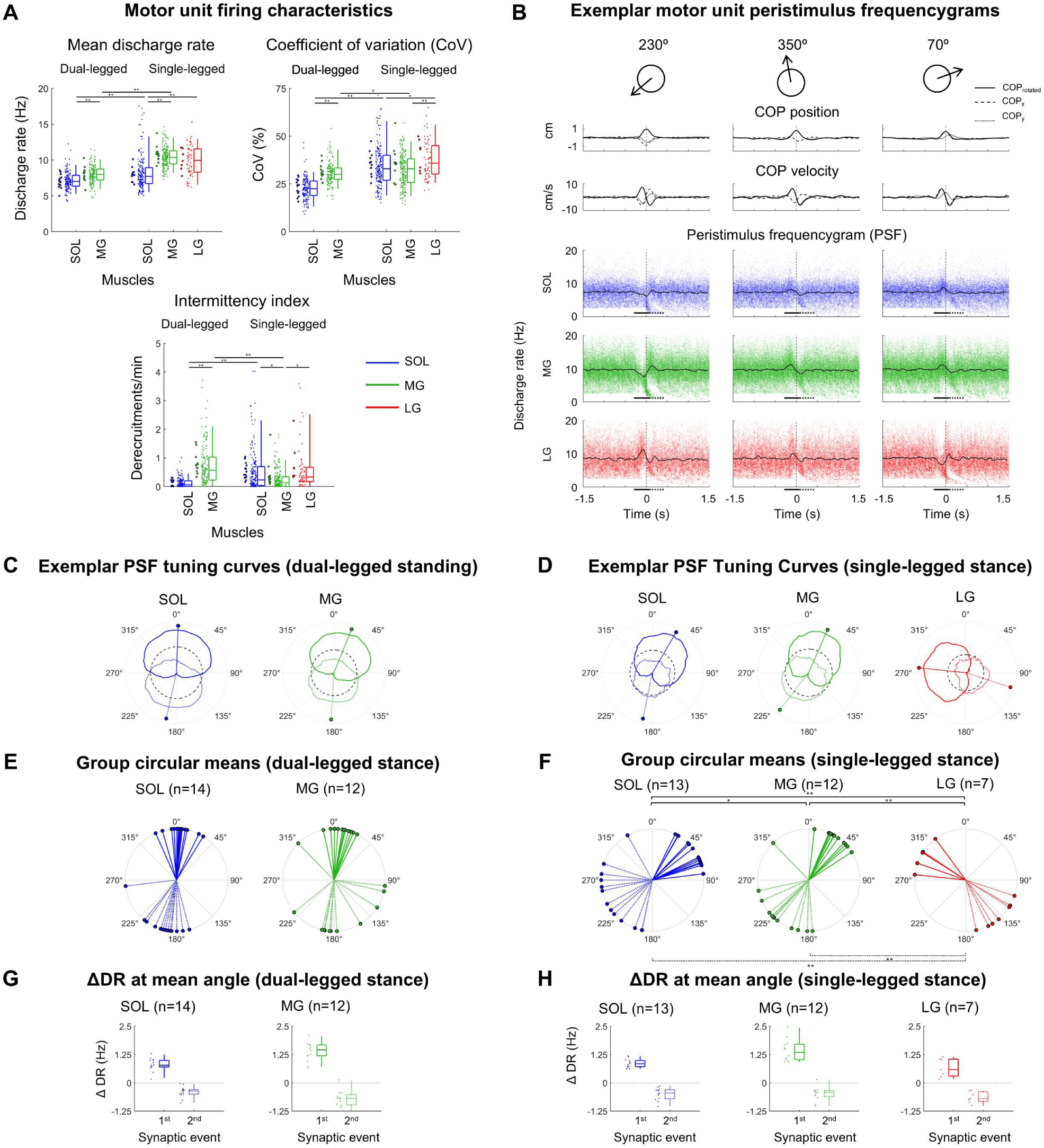
Motor unit characteristics and peristimulus frequencygram (PSF) tuning curves. (A) Motor unit discharge characteristics (mean discharge rate, coefficient of variation (CoV) of the interspike interval, and intermittency) across stance conditions and muscles. Box plots illustrate medians, 1^st^ and 4^th^ quartiles, and 5^th^ and 95^th^ percentiles (whiskers) for individual motor units (small data points) for the trial of each condition that had the highest motor unit yield from each participant. The leftmost data points represent individual participant means. Statistically significant Bonferroni-corrected pairwise comparisons for the effect of task and muscle (*p<0.05; **p<0.001) are shown above. (B) Exemplar PSFs are shown for COP events at three contrasting angles (230°, 350°, 70°) for an exemplar participant. On each PSF, individual data points represent the instantaneous discharge rate of single motor units; the black trace is a moving average of discharge rates. Upper traces illustrate waveform averaged COP position and velocity. Exemplar PSF tuning curves are shown for a single participant across both stance conditions (C,D). Solid lines represent tuning curves constructed from the initial synaptic period (-350 to 50 ms) while transparent lines represent those constructed from the secondary synaptic period (-50 to-450 ms; analysis windows are indicated by solid and dotted lines along the x-axis below each exemplar PSF (B)). Dashed, black radial values on each tuning curve represent a change in discharge rate of zero, such that values outside and within this region represent excitatory and inhibitory responses, respectively. PSF tuning curve circular means for the initial (solid line) and secondary (dotted line) synaptic periods across all participants with a sufficient number of motor units for each muscle are shown for both stance conditions (E,F).

This time-dependent antagonism between the muscles of the triceps surae is evident from examination of individual participant PSF tuning curves (Figure 3D). These illustrate that there are regions where muscles receive the same or opposite polarity of synaptic input depending on the orientation of COP movement. These also illustrate that there is clear directional tuning of both the initial and secondary synaptic event that straddle the peak of COP deflections in different directions. Circular mean angles of the initial synaptic event were consistent with our main analyses, with WW tests confirming significant differences in directional tuning between all three muscles during single-legged standing (SOL vs MG: F_(1,22)_=10.1182; p=0.0043; SOL vs LG: F_(1,12)_=141.5484; p<0.0001; MG vs LG: F_(1,12)_=132.3372; p<0.0001; Figure 3D, solid lines). The secondary synaptic event of opposite polarity also demonstrated directional tuning (Figure 3D, dotted lines), with WW tests confirming significant differences between SOL and LG (F_(1,12)_=39.0906; p<0.0001) and MG and LG (F_(1,12)_=43.9023; p<0.0001), but not SOL and MG (F_(1,22)_=1.3910; p<0.2508; Figure 3D, dotted lines). In addition to showing directional tuning, changes in motor unit firing rate during the secondary synaptic window at each muscle’s mean tuning angle were significantly different than 0 (all p-values <0.0001) and of opposite polarity relative to the initial synaptic event (Figure 3E).

These same analyses were applied to SOL and MG motor unit data from the dual-legged standing condition (Figure 3CEG). Consistent with the main analyses, PSF tuning curves for both muscles were oriented primarily along the sagittal plane, although mean angles for neither the initial and secondary synaptic events did not significantly differ between muscles (initial: F_(1,22)_=3.0278; p=0.0955; secondary: F_(1,22)_=1.7163; p=0.2037).

Watson-Williams tests indicating significant differences in circular means for the effect of muscle within each task are shown for the initial (above) and secondary (below) synaptic periods (*p<0.05; *p<0.001). (G and H) Show the change in discharge (ΔDR) of motor units during the initial (-350 to 50 ms) and secondary (-50 to 450 ms) synaptic periods at each muscle’s preferred angle of discharge (i.e., during COP deflections oriented at the mean angle of the initial synaptic event PSF tuning curve).

### Phasic activation of each muscle of the triceps surae results in heterogenous and task-specific COP trajectories

Peaks in the rectified and smoothed EMG were identified and used as event times to examine how each muscle of the triceps surae contributed to unique movement trajectories of the COP. Exemplar data presented in Figure 4 show these events separately for each muscle of the triceps surae during a single-legged standing trial alongside waveform averaged EMG and COP. These data illustrate that during single-legged standing, prominent phasic activation of the SOL were accompanied by corresponding activation of the MG, but not the LG (Figure 4B, top row). By contrast, during prominent phasic activation of the LG, neither the SOL nor MG modulated their activity (Figure 4B, bottom row). These effects were confirmed across the group data by calculating mean activity of each muscle’s waveform average within a ±250 ms peristimulus period centered around each EMG burst event (Figure 4E). During single-legged standing, 1-sample t-tests revealed that the mean response of the SOL and MG were not significantly different from zero during isolated activation of the LG (t_(12)_=-1.4147, p=0.1826; t_(12)_=-0.8177, p=0.4295, respectively; Figure 4E). Similarly, during phasic activation of either the SOL or MG, the mean response of the LG was also not different from zero (t_(12)_=-1.1531, p=0.2713; t_(12)_=0.2811, p=0.7834, respectively). By contrast, during phasic SOL activity, the mean response of the MG was substantially greater than zero (t_(12)_=11.3579; p<0.0001), and the same was true for the response of the SOL during bursts of MG activity (t_(12)_=6.3583; p<0.0001). This indicates that during periods where the LG is most active, the SOL and MG receive do not receive significant excitatory drive, and vice versa, while SOL and MG are more likely to be coactivate.

**Figure 4.**
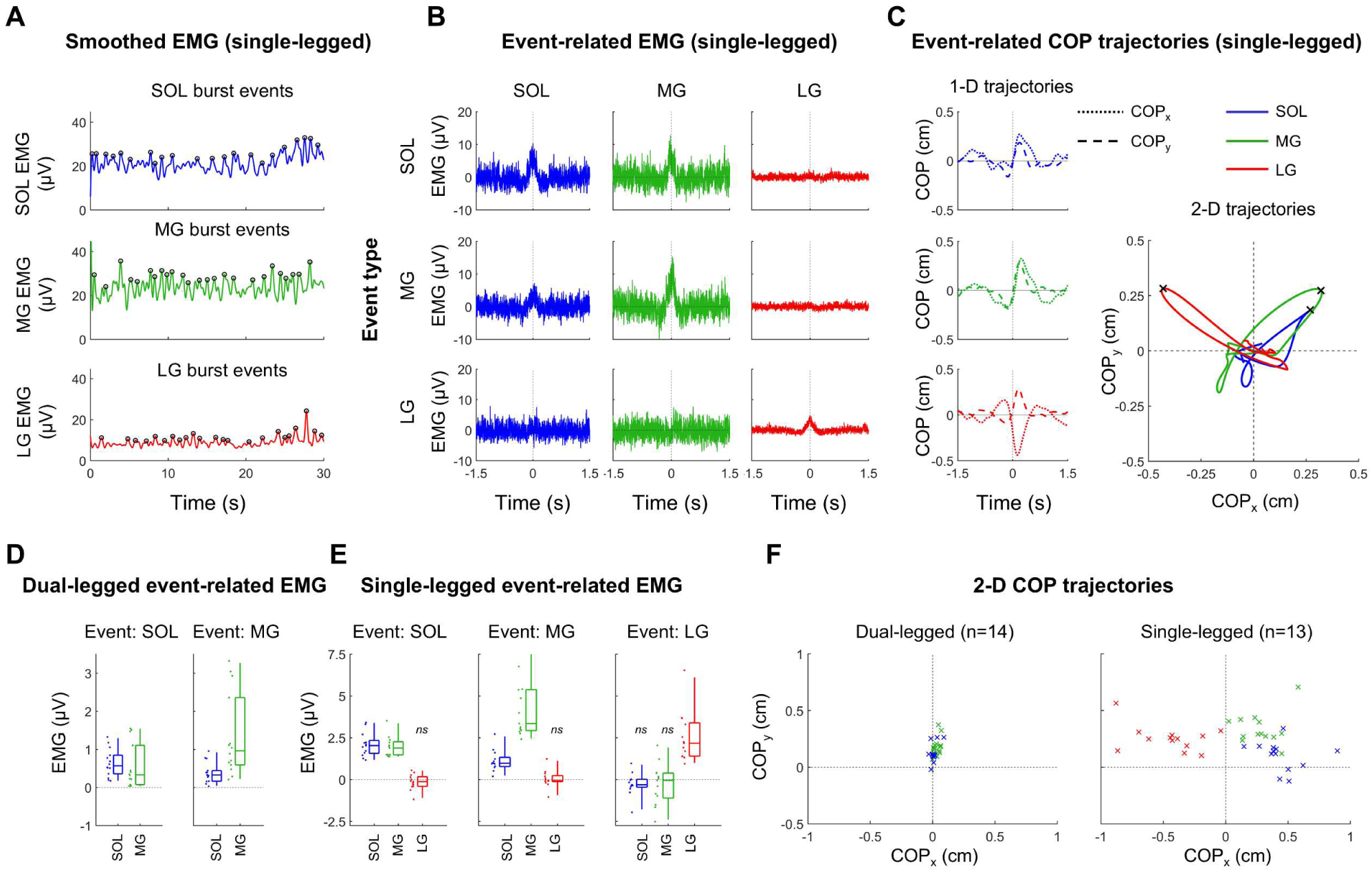
EMG burst analyses and 2-D COP trajectories. (A) Prominent bursts of activity from each muscle of the triceps surae were identified from rectified and smoothed (2 Hz low-pass) EMG (○). (B) Rectified EMG from each muscle was trigger-averaged to burst events from each muscle. From each of these waveforms, baseline-corrected mean rectified EMG between-250 ms and 250 ms was calculated (D, E). Note, the activity of all parts of the triceps surae are shown in response to the same type of EMG burst event, including the mean activity of the same muscle (e.g., LG activity during LG burst events). Single-sample t-tests were performed on mean event-related EMG, with non-significant effects (*ns*) indicating that a muscle did not modulate it activity during phasic activation of a given muscle (D, E). (C) COP_x_ and COP_y_ data were trigger-averaged to the same EMG burst events from each muscle. COP_x_ and COP_y_ waveform averages were combined and the apex of the 2-D COP trajectory was identified for each participant. 2-D COP trajectories in response to isolated activation of each muscle are shown for an exemplar participant during single-legged standing (C) with the apex of each trajectory labelled with an x. (F) Shows the apices of 2-D COP trajectories (color-coded x’s) for all participants during both stance conditions.

This independent activation pattern between the LG and SOL/MG likely owes to the differential effects each muscle has on the movement trajectory of the COP during single-legged standing. Figure 4C shows exemplar COP waveform-averaged relative to the same EMG burst events discussed above. These data reveal highly divergent trajectories of the COP, particularly along the frontal plane (Figure 4F). WW tests confirmed that the resultant angle of COP trajectories following phasic activation of each muscle significantly differed during single-legged standing (F_(2,36)_=152.8857, p<0.0001), with phasic activation of the LG resulting in more medially oriented COP trajectories compared to the SOL (F_(1,24)_=289.7355; p<0.0001) and MG (F_(1,24)_=167.8881; p<0.0001). While phasic activation of both the SOL and MG generally caused forward and lateral movement of the COP, WW tests indicated that their movement trajectories were also significantly different, with phasic activation of the SOL resulting in more laterally oriented COP trajectories than the MG (F_(1,24)_= 23.1603; p<0.0001; Figure 4F). No significant differences in the resultant distance of COP trajectories following prominent phasic activation of each muscle were observed (F_(2,24)_=1.205, p=0.317).

When these same analyses were applied to the data from the dual-legged standing condition, prominent phasic activation of the SOL was found to be accompanied by significant activation of the MG (t_(13)_=3.5848; p=0.0033), and vice versa (t_(13)_=5.3515; p<0.0001; Figure 4D). Likewise, both muscles caused similar COP movement trajectories oriented primarily along the sagittal plane (F_(1,26)_=0.9278, p=0.3443), although activation of the MG caused a larger resultant displacement of the COP (t_(13)_=3.618, p=0.0031; Figure 4F; Table S1).

### Within-muscle directional tuning is homogenous

To determine if the stark between-muscle differences in directional tuning observed during single-legged standing were further refined within each muscle, PSTH tuning curves were constructed from regionally-defined motor unit subpopulations (Figure 5AB). PSTH tuning curves could be constructed from groups of motor unit spike trains (≥4 units per group) with MUAP centroids sufficiently separated along the x-(mediolateral) and y-axes (proximodistal) for the SOL (medial vs lateral: n=10, Δ x-centroid location=19.3 ±4.4 mm; proximal vs distal: n=11; Δ y-centroid location=51.6 ±8.8 mm), MG (medial vs lateral: n=8, Δ x-centroid location=16.2 ±3.4 mm; proximal vs distal: n=7; Δ y-centroid location=44.7 ±8.6 mm), and LG (medial vs lateral: n=5, Δ x-centroid location=19.1 ±4.6 mm; proximal vs distal: n=5; Δ y-centroid location=52.0 ±14.3 mm). WW tests reveal no significant difference in mean tuning angles within muscles for either medial vs lateral comparisons (SOL: F_(1,18)_=1.9385, p=0.1808; MG: F_(1,14)_=0.0104, p=0.9204; LG: F_(1,8)_=0.6332, p=0.4492; Figure 5C) or proximal vs distal comparisons (SOL: F_(1,20)_=0.4523, p=0.5089; MG: F_(1,12)_=0.0076, p=0.9318; LG: F_(1,8)_=0.0971, p=0.7634; Figure 5D). This within muscle homogeneity of directional tuning orientation is consistent with supplemental analyses demonstrating minimal variability in tuning curve angles constructed from all differential electrode pairs spanning the surface of the electrode array (Figure S2).

**Figure 5.**
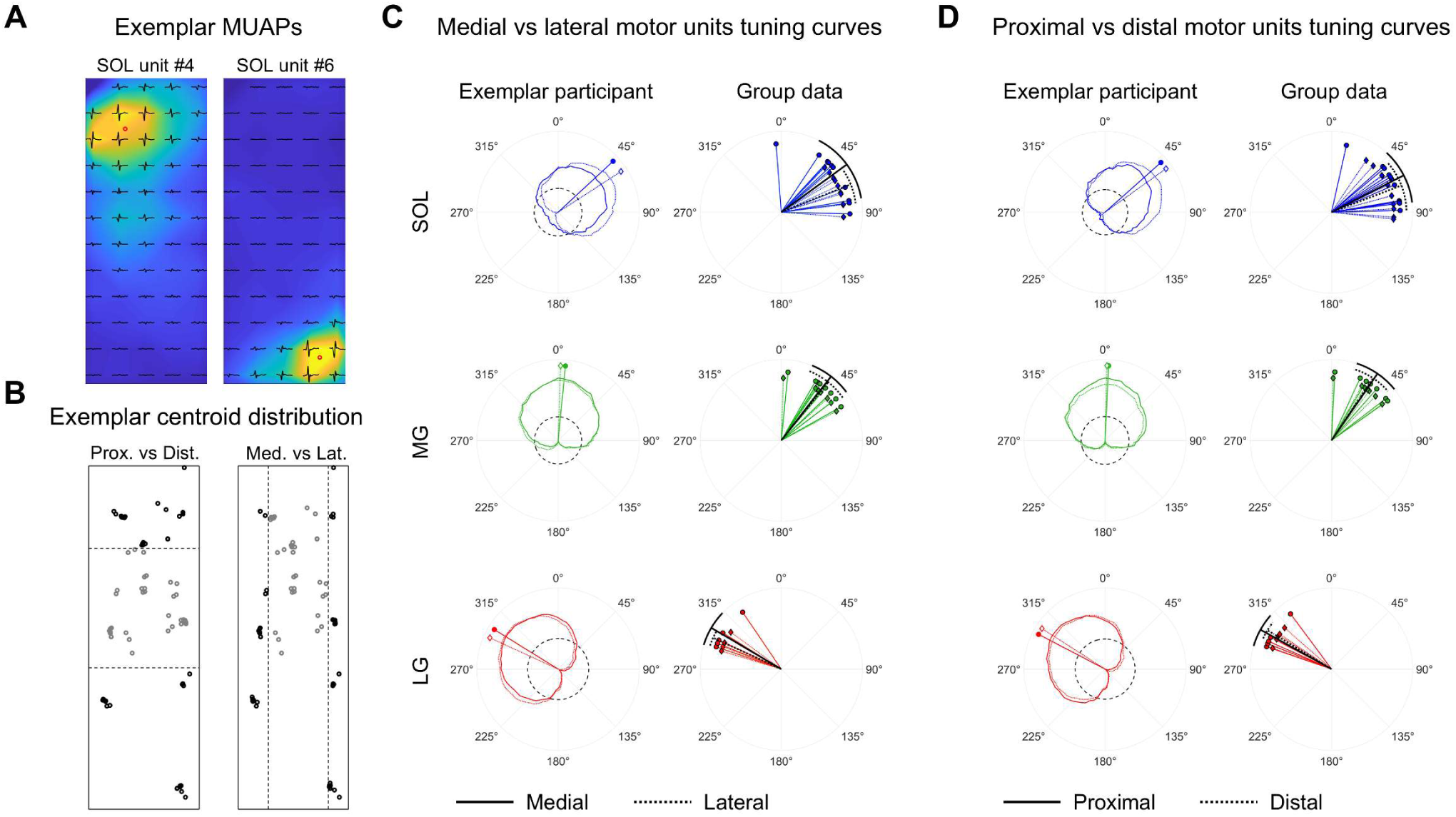
Tuning curves constructed from regionally defined motor unit subpopulations during single-legged standing. (A) Motor units action potentials (MUAPs) were mapped onto recording arrays for each muscle using spike-triggered averaging procedures and the xy coordinates of the centroid were identified (red circle on exemplar MUAP heat plots). (B) Illustrates all MUAP centroids across 8 trials of single-legged standing for an exemplar participant (slight jitter was applied to each centroid location for illustrative purposes only). Motor units were grouped according to centroid location using quartile splits (boundaries shown as dashed lines). Black dots denote motor unit centroids that fell outside of respective proximal/distal and medial/lateral quartile boundaries, which were then used to construct regional PSTH tuning curves for all participants that had a sufficient yield of motor units. (C,D) Show exemplar regional PSTH tuning curves for medial (solid) vs. lateral (dotted) and proximal (solid) vs. distal (dotted) units as well as group data showing all circular means for the same comparisons. Grand circular means and standard deviations are superimposed on all plots (black solid and dotted lines with outer radial bands).

Mediolateral regional differences in the SOL were examined with greater spatial contrast in a subset of participants where recordings were made from separate arrays placed over the medial and lateral SOL. Motor units from the lateral array were classified as either ‘lateral SOL’ or ‘putative fibularis’, while those from the medial array were classified as medial SOL (Figure 6B). While PSTH tuning curves for medial and lateral SOL units were virtually identical (F_(1,16)_=0.0078, p=0.9308), putative fibularis units showed near opposite directional tuning to both medial (F_(1,10)_=60.2928, p<0.0001) and lateral (F_(1,10)_=56.4824, p<0.0001) SOL units. That is, unlike both groups of SOL units, which demonstrated the greatest discharge probability prior to forward and lateral COP peaks, putative fibularis units demonstrated the greatest discharge probability prior to medial COP movement (Figure 6D). Linear mixed effects models revealed that motor unit firing and morphological characteristics differed significantly amongst these classifications of motor units (discharge rate: F_(2,179)_=58.531, p<0.0001; CoV: F_(2,179)_= 17.837, p<0.0001; intermittency: F_(2,180)_=41.979, p<0.0001; MUAP spike height: F_(2,183)_=15.629, p<0.0001). Follow-up Bonferroni-corrected pairwise comparisons revealed that in all cases, these characteristics did not significantly differ between medial and lateral SOL units (p-value range: 0.0713-0.9985), but instead between putative fibularis units and both classifications of SOL units. In particular, putative fibularis units demonstrated greater discharge rates (fibularis vs medial SOL: t_(180)_=10.150, p<0.0001; fibularis vs lateral SOL: t_(181)_=10.275, p<0.0001), CoV (fibularis vs medial SOL: t_(180)_=5.793, p<0.0001; fibularis vs lateral SOL: t_(182)_=5.453, p<0.0001), intermittency (fibularis vs medial SOL: t_(181)_=8.867, p<0.0001; fibularis vs lateral SOL: t_(182)_=8.372, p<0.0001), and MUAP spike heights (fibularis vs medial SOL: t_(183)_=4.422, p<0.0001; fibularis vs lateral SOL: t_(184)_=5.590, p<0.0001; Figure 6C).

**Figure 6.**
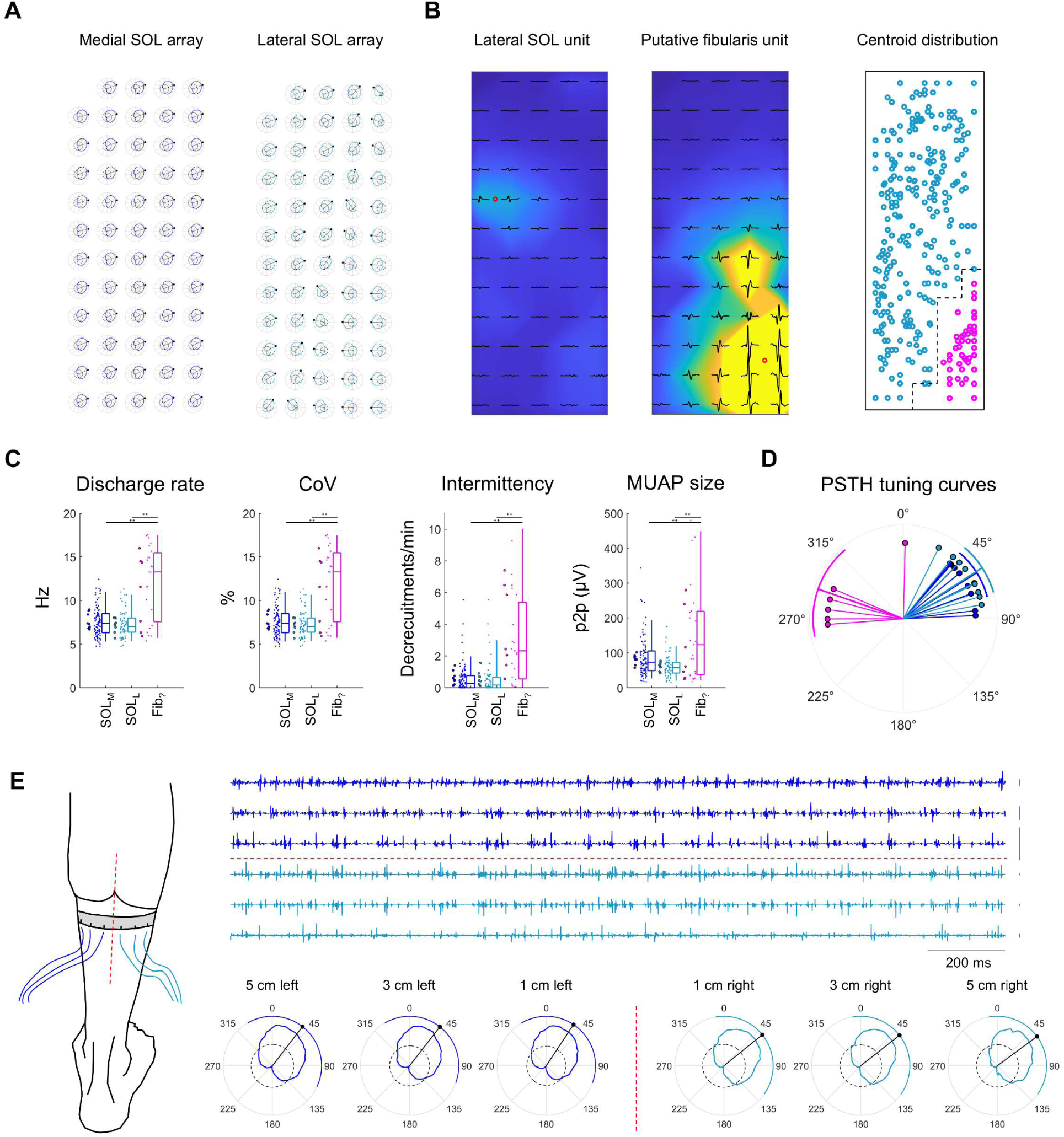
Medial vs lateral SOL tuning curve and motor unit analyses during single-legged standing. (A) Tuning maps from the medial (blue) and lateral (turquoise) SOL electrode arrays are shown for an exemplar participant. (B) Motor units decomposed across the lateral SOL array were grouped according to centroid location as either lateral SOL (turquoise) or putative fibularis (pink). The centroid of all motor units from the lateral SOL array for all participants, along with the boundaries used to classify them, are shown on the rightmost plot of (B). (C) Box plots show measures of motor unit discharge characteristics and morphology for each unit classification, including units from the medial SOL array (deep blue). Statistically significant pairwise differences between motor unit classifications (*p<0.05; **p<0.001) are shown above data. Peristimulus-time histogram (PSTH) tuning curves were constructed for each classification when there were sufficient number of decomposed motor units. (D) Circular means were calculated from each tuning curve and are plotted alongside color-coded grand circular means and standard deviations. (E) In a separate experiment, 6 pairs of fine wire electrodes were inserted into the SOL of the right leg in a male participant (33 years old) who performed multiple trials of single-legged standing. Electrodes were separated by 2 cm increments and spanned the entire mediolateral width of the SOL. Tuning curves were constructed from the rectified EMG at each location using the same procedures outlined in Figure 1.

To further examine mediolateral regional differences in SOL directional tuning without introducing cross-talk from adjacent muscles, tuning curves were constructed from fine wire EMG recordings spanning the entire mediolateral margins of the SOL in 2 cm increments in one male participant (Figure 6E). These tuning curves were largely consistent and did not demonstrate any clear mediolateral gradation (Figure 6E).

## Discussion

This study demonstrates that each triceps surae muscle can generate different torques about the ankle, and the nervous system exploits this functional heterogeneity to tune balance corrective adjustments in a task-dependent manner. This was shown through convergent analyses that examined muscle activity and COP movements during two balance tasks that use distinct neuromuscular control strategies to maintain equilibrium.

The principal analyses of this study examined muscle activity time-locked to prominent peaks in unidimensional COP time-series rotated across 360°. Reversals in COP position formed the basis of these analyses because they coincide with prominent changes in neural drive to the muscles generating balance corrective torques (Loram et al. 2002; 2005; Zaback et al. 2023). This made it possible to construct EMG tuning curves without the need for multi-directional perturbations (Safavynia and Ting 2013; Monte et al. 2024). Consistent with previous work, only the SOL and MG were active during dual-legged standing, and each modulated their activity uniformly with COP deflections along the sagittal plane (Heroux et al. 2014). By contrast, when standing on one leg, all three muscles of the triceps surae showed robust activation and highly divergent directional tuning. Tuning curves for the SOL and MG were rotated clockwise relative to dual-legged standing, with SOL tuning curves oriented more laterally than MG, indicating that each muscle was most active during plantarflexion-*inversion* balance corrective torques, with the SOL having a greater inversion component. The LG showed an almost opposite pattern of activity, being most active during plantarflexion-*eversion* balance corrective torques. These task-and muscle-specific differences were confirmed by motor unit PSF tuning curves, which demonstrated that motor units from each triceps surae muscle could show the same or opposite change in firing rate relative to one another depending on the orientation of the balance corrective torque.

While these findings are consistent with previous work that has suggested that each triceps surae muscle receives largely independent neural input (Hug et al. 2021; Rossato et al. 2024), they provide a functional explanation for why this independent control is advantageous. This is clearly illustrated through the analyses of 2-D COP trajectories time-locked to prominent bursts of EMG, which demonstrate how independent activation of each muscle causes movement of the COP in different directions during single-legged standing. These observations are congruent with anatomical features of the triceps surae. The tendonous bundles (subtendons) from each triceps surae muscle twist as they descend and fuse into the Achilles tendon, with each subtendon inserting onto a separate region of the calcaneal tuberosity (Szaro et al. 2009; Edama et al. 2015; 2016; Lehr et al. 2021). This partitioned organization allows each muscle to generate distinct torques at the ankle joint (Arndt et al. 1999). *In vitro* stimulation of each triceps surae muscle in human cadavers has shown that while all three generate a plantarflexion torque, the LG generates an eversion torque, while the SOL and MG generate inversion torques (Arndt et al. 1999). While this corresponds with what we observed *in vivo* during single-legged standing, the SOL and MG appeared to generate near pure plantarflexion torques during dual-legged standing.

This likely owes to the strong mechanical coupling between the ankles and hips during dual-legged standing (Day et al. 1993). This coupling reduces the amount of inversion-eversion that occurs at the ankle joint, constraining the SOL and MG to act as plantar flexors during this task (Winter et al. 1993). Only once there is little mechanical coupling between the ankles and hip, as is the case during single-legged standing, do their distinct actions along the frontal plane become apparent. This is not to say that triceps surae do not have unique roles during dual-legged standing. Differences in motor unit firing characteristics (Heroux et al. 2014) and muscle fibre architecture and composition (Edgerton et al., 1975; Friederich & Brand 1990; Johnson et al., 1973) suggest the SOL may be well-suited for controlling low-frequency sway, while the MG is better for generating larger and more rapid adjustments (Dakin et al. 2016). This is consistent with the observation that phasic activation of the MG propelled the COP farther than the SOL during dual-legged standing. Thus, independent control of the triceps surae may still be advantageous during tasks where each muscle is constrained to generate similarly oriented balance corrective torques.

By examining changes in motor unit firing rates, we observed that COP reversals are often straddled by two synaptic events of opposite polarity. The nature of this pattern of neural drive is unclear, but can be speculated based on the underlying neuromechanical control processes. Each reversal marks a moment when the COP has moved to corral the center of mass, either to slow its movement in a particular direction or reverse it entirely. These adjustments may reflect open-loop, predictive control processes shaped by intermittent, rather than continuous feedback (Gawthrop et al. 2014; Loram et al. 2022). That is, when sway or a predicted sensory state exceeds an instability threshold, an intermittent correction is triggered to restore equilibrium (Loram et al. 2006; Bottaro et al. 2008). It follows that the first synaptic event reflects the primary neural drive to generate the balance corrective adjustment that is predicted to restore stability. The nature of the secondary synaptic event is less clear. It may reflect a preprogrammed braking mechanism to dampen the initial balance correction, permitting a rapid response with minimal sway oscillation, similar to that seen during ballistic reaching movements (Hallet et al. 1975). Alternatively, it could reflect a separate, event-driven response to reafferent signals indicating a prediction error with the initial balance correction (Forbes et al. 2018). Regardless of the nature of this secondary synaptic event, it demonstrated directional tuning opposite to the first event, indicating there is coordination of neural drive both within and between muscles to generate balance corrective actions.

While each triceps surae muscles generates distinct balance corrective torques during single-legged standing, this heterogeneity could be refined by distinct motor unit subpopulations or task groups with each muscle (Loeb 1985; Siegal, 1992; Vieira et al. 2011). Such intramuscular differences have been speculated based on known anatomical compartments with distinct fibre architectures and innervation from separate muscle nerve branches (Segal et al. 1991; Loh et al. 2003; Ashalou et al. 2014; Bolsterlee et al. 2018). Furthermore, recent work suggests that separate motor unit populations within each triceps surae muscle may be independently recruited to accommodate different postural orientations (Cohen et al. 2021; 2023). Despite this, we found no evidence of functionally distinct motor unit task groups within any muscle of the triceps surae. PSTH tuning curves from regionally-defined motor unit subpopulations were highly similar within each muscle and there was minimal variability in EMG tuning curve orientation when constructed from differential electrode pairs spanning the surface of each array (Figure S2). Thus, while functional subdivisions may exist, our data provide little evidence that nervous system leverages these to tune balance corrective torques.

While these regional analyses were limited by a relatively narrow recording volume along each muscle’s mediolateral axis, it was possible to expand this for the SOL in a subset of participants where recordings were made from separate medial and lateral arrays. Consistent with work demonstrating that neural drive is largely shared between medial and lateral portions of the SOL (Aeles et al. 2023), directional tuning was virtually identical for medial and lateral SOL units and a separate fine wire experiment showed no mediolateral gradation in tuning curve orientation.

An inadvertent finding from these analyses was the presence of a motor unit population likely originating from another muscle rather than separate compartment within the SOL. These units were localized to the distal-lateral corner of the lateral SOL array, consistent with proximity to the fibularis muscles, and showed markedly different firing characteristics from both medial and lateral SOL units. Tuning curves for these putative fibularis units were diametrically opposed to both groups of SOL units, consistent with the fibularis muscles’ role in eversion (Mendez-Rebolledo et al. 2021; Kunugi et al. 2023). MUAPs for putative fibularis units were often large and contaminated large portions of the lateral SOL array, suggesting that limiting decomposition to the anatomical borders of a muscle may be insufficient in eliminating crosstalk. Instead, oversampling EMG data from surrounding muscles and assigning motor units after decomposition and MUAP reconstruction may more effectively minimize cross-talk.

This study demonstrates that the activity of each muscle of the triceps surae can be independently tuned to generate balance corrective adjustments in a task-dependent manner. This fractionated control of closely related lower limb postural muscles is likely critical to the maintenance of equilibrium and may be impaired by various musculoskeletal (e.g., chronic ankle instability) and neurological (e.g., stroke) impairments. Future work will investigate task-dependent tuning across a variety of lower limb postural muscles in healthy and pathological populations to improve our understanding of the causes of functional balance deficits and potentially develop targeted biomarkers of impaired neural control.

## Additional Information

## Data Availability

All data generated as part of this experiment will be made available upon reasonable request.

## Acknowledgements

The authors acknowledge Matt Topley for his technical assistance.

## Software Accessibility

All code used to process data and generate figures will be made available upon reasonable request.

## Author Contributions

This study was conducted in Professor Christopher K. Thompson’s lab at Temple University in Philadelphia, PA, USA. M.Z. contributed to the conception and design of the study, data acquisition, analysis, interpretation of data, and drafting and revising the manuscript. C.K.T. contributed to the conception and design of the study, interpretation of the data, and drafting and revising the manuscript. All authors approved the final version of the manuscript and agree to be accountable for all aspects of the work in ensuring that questions related to the accuracy or integrity of any part of the work are appropriately investigated and resolved. All persons designated as authors qualify for authorship, and all those who qualify for authorship are listed.

## Appendix

**Figure S1.**
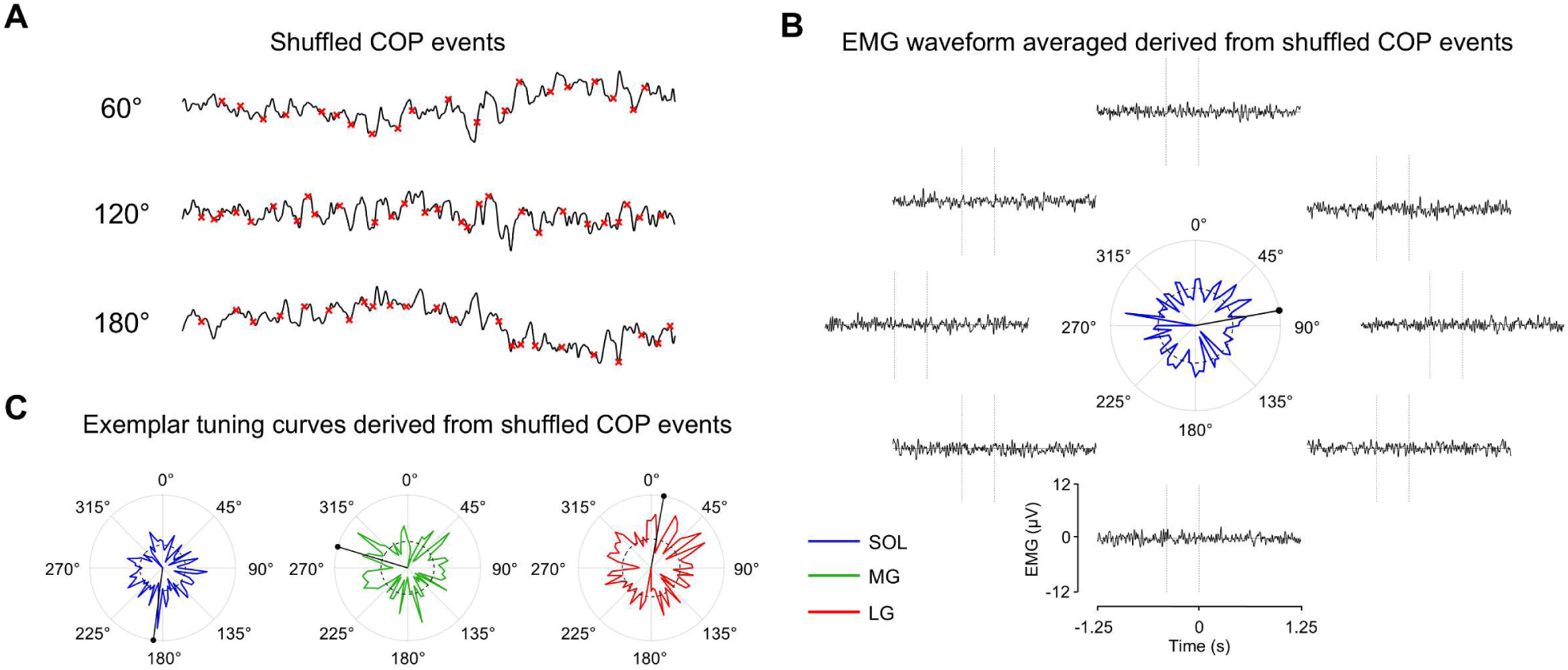
Validation of tuning curve analyses. For each iteration of COP rotation, event times were shuffled, such that they had the same frequency of occurrence, but no relation to the identified postural events as illustrated for 3 exemplar rotated COP time-series (A). (B) Illustrates the reconstruction of a SOL EMG tuning curve from shuffled COP events during single-legged standing for an exemplar participant. No visible modulation of EMG was observed at any iteration of COP rotation (B) nor were any non-circular distributions or regions of continuous excitatory/inhibitory responses for any muscle of the triceps surae (C).

**Figure S2.**
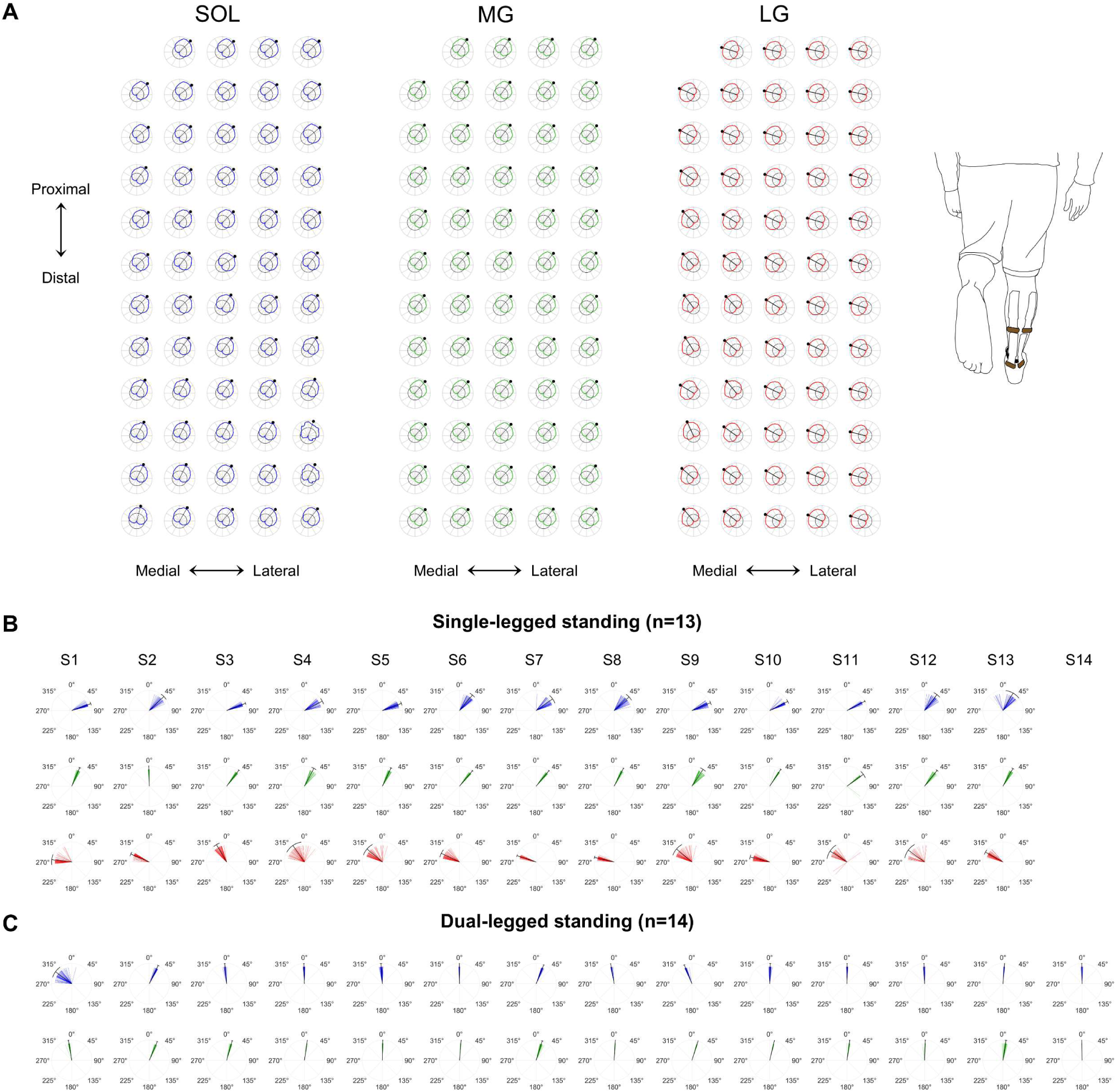
EMG tuning maps. (A) For each muscle, EMG tuning curves were constructed for each differential pair of vertically-adjacent electrodes plotted according to their physical location on the array. All circular means from each tuning curve spanning each array were plotted on polar coordinates separately for each participant during single-(B) and dual-legged (C) standing. The grand circular mean is superimposed as a solid black line on each polar plot and the circular standard deviation of all tuning curves is plotted as an outer radial band.

---

**Table S1:** Descriptive statistics for EMG and PSF tuning curve, motor unit firing characteristics, and 2-D COP trajectory analyses.

|  | Dual-legged standing |  | Single-legged standing |  |  |
| --- | --- | --- | --- | --- | --- |
|  | SOL | MG | SOL | MG | LG |
| EMG tuning curves |  |  |  |  |  |
| Mean angles (°) | 1.7±17.7 | 12.5±8.6 | 61.0±14.5 | 36.1±13.1 | 303.0±16.6 |
| Motor unit characteristics |  |  |  |  |  |
| Motor unit # (per trial) | 8.7±4.1 | 7.4±7.7 | 12.5±8.2 | 13.6±10.2 | 4.0±5.3 |
| Discharge rate (Hz) | 7.2±1.4 | 7.8±1.2 | 8.1±2.2 | 10.4±1.5 | 9.7±1.9 |
| CoV of ISI (%) | 22.6±6.5 | 31.8±6.0 | 35.0±11.7 | 33.0±8.8 | 38.1±10.0 |
| Intermittency | 0.2±0.3 | 1.0±0.9 | 0.5±0.8 | 0.3±0.4 | 0.6±0.7 |
| Motor unit PSF tuning curves |  |  |  |  |  |
| 1<sup>st</sup> event angle (°) | 3.8±13.7 | 10.1±10.7 | 55.7±17.9 | 32.3±13.3 | 300.0±13.4 |
| 2<sup>nd</sup> event angle (°) | 196.0±22.3 | 171.3±48.7 | 244.2±37.5 | 221.5±34.1 | 137.2±19.7 |
| 2-COP trajectories |  |  |  |  |  |
| Trajectory angle (°) | 2.0±30.7 | 10.6±7.8 | 74.3±18.4 | 38.4±17.7 | 302.7±17.2 |
| Trajectory dist. (mm) | 1.3±0.8 | 1.9±0.8 | 4.8±1.6 | 4.6±1.5 | 5.3±2.4 |

Means and standard deviations are provided for circular mean angles derived tuning curves constructed from rectified EMG (EMG tuning curves) and motor unit PSFs. For motor unit PSFs, separate tuning curves were constructed from changes in firing rate before (1<sup>st</sup> event: -350 to +50 ms) and after (2<sup>nd</sup> event: +50 ms to +450 ms) peak COP deflections throughout 2D space. Motor unit discharge characteristics are provided for all motor units decomposed across all 8 dual- and single-legged trials.

